# Symbiotic bacterial community of *Drosophila melanogaster* changes with nutritional modifications of the diet but can alleviate negative effects on larval phenotypes

**DOI:** 10.1101/2021.09.07.458894

**Authors:** Andrei Bombin, Owen Cunneely, Sergei Bombin, Kira Eickman, Abigail Ruesy, Rachael Cowan, Abigail Myers, Mengting Su, Jonathan Mosley, Jane Ferguson, Laura Reed

**Affiliations:** Vanderbilt University Medical Center; The University of Alabama; Yale University

## Abstract

Obesity is an increasing pandemic and is caused by multiple factors including genotype, psychological stress, and gut microbiota. Our project investigated the effects produced by high fat and high sugar dietary modifications on microbiota and metabolic phenotype of *Drosophila melanogaster*. Larvae raised on the high fat and high sugar diets exhibited bacterial communities that were compositionally and phylogenetically different from bacterial communities of the larvae raised on normal diets, especially if parental microbiota were removed. Several of the dominant bacteria taxa that are commonly associated with high fat and high sugar diets across model organisms and even human populations showed similar pattern in our results. *Corynebacteriaceae* and *Erysipelotrichaceae* were connected with high fat food, while *Enterobacteriaceae* and *Lactobacillaceae* were associated with high sugar diets. In addition, we observed that presence of symbiotic microbiota often mitigated the effect that harmful dietary modifications produced on larvae, including elevated triglyceride concentrations and was crucial for *Drosophila* survival, especially on high sugar peach diets.

## INTRODUCTION

The pandemic of obesity contributes to the increase in multiple health conditions including hypertension and type two diabetes mellitus among many others (Seganfredo et al., 2017, Wahba and Mak, 2007, Thompson et al., 2007). The problem of obesity is further complicated by the diversity of causal factors such as dietary habits, genotype, epigenetic regulation, psychological stress, sleep deprivation, and gut microbiota composition (Seganfredo et al., 2017, Han and Lean, 2016). These issues make the development of an efficient treatment a complex problem requiring a multivariate research approach (Thompson et al., 2007, Kaur, 2014).

Diet influences metabolic phenotypes and longevity of *Drosophila*. Raising *D. melanogaster* larvae on a high sugar diet (86.4% of total calories available from carbohydrates) resulted in a smaller body size, delayed development, elevated glucose, trehalose, and reduced glycogen concentrations of the larvae (Musselman et al., 2011). In addition, those larvae developed a reduced insulin sensitivity and had higher triglyceride concentrations, overall exhibiting a diabetic phenotype (Musselman et al., 2011). A high sugar diet also significantly decreases the pupal weight (Reed et al., 2010). Obesity, induced by high sugar diet, increases with aging of the flies (Skorupa et al., 2008). A high fat diet can also produce an obese phenotype in *Drosophila*, which results in higher triglyceride levels and disturbs insulin homeostasis (Birse et al., 2010). In addition, a diet rich in fats leads to similar effects in body mass and development rate as a high sugar diet and also produces transgenerational effects (Musselman et al., 2011, Dew-Budd et al., 2016). An obese phenotype is harmful for *Drosophila* fitness. Higher fat mass storage was shown to reduce longevity of flies and resulted in lower fecundity (Moghadam et al., 2015). On the opposite side of the spectrum, diets restricted in sugars and/or yeast consumption were shown to increase life span (Mair et al., 2005).

In *D. melanogaster*, microbiota influence several life history traits and metabolic phenotypes such as survival untill pupation, development time, weight, protein, triglyceride, and glucose concentrations (Newell and Douglas, 2014, Dobson et al., 2015, Huang and Douglas, 2015). Interestingly, the impact of harmful diets, such as high fat and high sugar, forms a phenotype similar to flies that have a removal of symbiotic microbiota. For example, axenic flies often exhibit elevated glucose and triglyceride levels (Wong et al., 2014, Henry et al., 2020a). In addition, certain microbial taxa from *Acetobacter* and *Lactobacillus* genera may reduce host appetite for essential amino acids and increase the appetite for sugar consumption; thereby potentially providing the host with the essential amino acids and competing for available sugars (Leitão-Gonçalves et al., 2017).

Several studies demonstrated the differences in the symbiotic microbiota community of the wild flies and lab raised flies (Chandler et al., 2011, Vacchini et al., 2017, Tefit et al., 2017). In our previous work, we also observed that larvae raised on the natural peach diet and standard lab diet exhibited a substantial difference in the microbiota composition. It was also demonstrated that in *Drosophila* and mouse models, the effect that symbiotic microbiota provides on host phenotype may vary with diet, genotype, and their interactive effects (Wong et al., 2014, Kreznar et al., 2017, Henry et al., 2020a, Zhang et al., 2009, Org et al., 2015).

We raised *Drosophila* from five Drosophila Genetic Reference Panel (DGRP2) lines until late third instar larval stage on standard lab diets and peach-based diets modifying them with addition of extra sugars or fats, to mimic the westernized diet. In order to control for the action of live environmental and parental microbiota we also used an autoclaved variant of each of the peach-based diets and used larvae from sterilized and non-sterilized embryos. Therefore, our study is equipped to evaluate the interaction between harmful dietary modifications and *Drosophila*’s natural microbiota composition, while controlling for effects produced by genotype and parental microbiota. We hypothesized that nutritional modifications will cause shifts in larval bacterial community composition and metabolic phenotypes, however natural microbiota would alleviate negative metabolic effects caused by high fat and high sugar diets.

## MATERIALS AND METHODS

### Diet Preparation

Mixtures of naturally decayed peaches were prepared (homogenized) and stored, and then used to create the peach-based diet (peach regular - PR) using the procedures described in Bombin et al. (Bombin et al., 2020). The standard lab diet (R) diets were also prepared according to the protocols described previously (Dew-Budd et al., 2016, Mendez et al., 2016, Watanabe and Riddle, 2017, Bombin et al., 2020). In addition, high sugar and high fat diet variants were prepared for both peach-based diet and regular *Drosophila* lab-based diets as described below. All diet types were distributed with 10 mL of food per *Drosophila* culture vial.

To prepare the peach high-fat diet (PHF) we allowed one liter of the regular peach diet to thaw before supplementing it with coconut oil in an amount equal to 3% of the total mixture weight, then warmed the mixture to 28 °C (to melt the coconut oil). The mixture was then completely homogenized it with an immersion blender before distributing it into vials. To prepare the regular high-fat diet (RHF) we supplemented one liter of liquid *Drosophila* cornmeal-molasses standard lab diet with 30g of coconut oil (3%) during cooking, thoroughly mixed it and distributed it into vials to solidify. To prepare the peach high-sugar diet (PHS) we allowed one liter of the regular peach diet to thaw before supplementing it with sucrose in an amount equal to 11% by weight, then completely homogenized it with an immersion blender before distributing it into vials. In order to prepare peach high fat autoclaved (PHFA), peach high sugar 11% autoclaved (PHSA11), and peach high sugar 6% autoclaved (PHSA6) diets, vials containing the corresponding food were autoclaved for 25 min at 121 °C. Since our first results indicated that most of the tested genotypes could not survive on PHSA11 if the embryos went through a sterilization treatment, the PHSA6 diet was introduced in the second and third rounds of the experiment. PHSA6 diet was prepared as described above for PHSA11 but with only 6% extra sucrose added by weight. To prepare the regular high-sugar diet (RHS) we supplemented one liter of the *Drosophila* standard lab diet described above with 90 g of sucrose (to make 13% sugar diet which is similar in sugar content with our PHS diet) before the mixture solidified during cooking, then it was completely homogenized by thoroughly mixing it before distributing it into vials.

### Drosophila stocks and husbandry

For the high-sugar portion of the study, the following five naturally derived genetic lines were sourced from the Drosophila Genetic Reference Panel (DGRP2): **153, 748, 787, 802**, and **805** (Mackay et al., 2012, Huang et al., 2014). For the high-fat portion of the study, the following 10 naturally derived genetic lines were sourced from the DGRP2 project: **142, 153, 440, 748, 787, 801, 802, 805, 861**, and **882** (Mackay et al., 2012, Huang et al., 2014). These lines were selected because they showed a diversity of phenotypic responses to diet modifications in preliminary studies. These stocks were maintained using the procedures as described in previous works, maintaining them at a constant temperature (25 °C), humidity (50%), 12:12h light: dark cycle (Dew-Budd et al., 2016, Mendez et al., 2016).

### Drosophila embryos sterilization

In order to test for the effects of the parental microbiota, surface sterilized (S) and non-sterilized (NS) embryos were prepared as described in Bombin et al. (Bombin et al., 2020), prior to being placed on their respective diet treatments.

### Larval rearing and collection

Due to the nature of rearing and collecting and the large number of larvae required, this portion of the study was carried out in separate high-fat and high-sugar portions. Each portion of the study followed the same pattern. Across three separate time periods (∼25-30 days apart), 50 sterilized or non-sterilized larvae of each genetic line were added to each diet types. For the high-fat study, there were at least three vials each of PHF, PHFA, RHF, PA, PR and R diet. For the high-sugar component, there were at least three vials each of PHS, PHSA11 (from the second round, PHSA6% and PHSA11% to help mitigate low survival on a diet containing 11% sucrose), RHS, PA, PR, and R diets. For the HS study, only NS larvae were used for PR and R diets as controls. Larvae were allowed to develop until the late 3^rd^ instar stage and were subsequently collected, sorted, and stored frozen following the protocols described in Bombin et al. (Bombin et al., 2020).

### Measuring Experimental Phenotypes

The measured experimental phenotypes of **survival** (proportion of larvae survival to 3rd instar), **developmental rate** (days to 3rd instar larva), and **weight** of individual 3rd instar larvae were measured as described in Bombin et al. (Bombin et al., 2020). **Triglyceride** concentrations were quantified, by homogenizing 10 3^rd^ instar larvae per sample and measuring the total triglyceride concentration using the Sigma triglyceride Determination Kit (Clark and Keith, 1988, De Luca et al., 2005, Reed et al., 2010). Results were adjusted to represent the average triglyceride concentration per mg of dry larval weight. **Protein** concentrations were quantified using the Bradford’s method with 10 homogenized larvae per sample (with the exception of 3% of HF and 1% of HS samples, in which we used one to nine larvae due to especially low survival rates of the specific groups) (Bradford, 1976, Dew-Budd et al., 2016). Protein values were averaged to represent the protein concentration per mg of dry larvae weight. **Glucose** concentrations were quantified via homogenization of 10 larvae (with the exception of several HF and HS samples in which we used one to nine larvae) with subsequent overnight incubation in 1 μg/mL trehalase solution and further application of the Sigma Glucose Determination Kit as described in Bombin et al. (Bombin et al., 2020). Glucose concentrations were averaged and adjusted to represent the amount of glucose per mg of dry larval weight.

### DNA extraction and sequencing

Five genetic lines **153, 748, 787, 802**, and **805** raised on HF and HS diets were used for bacterial DNA 16S rRNA sequencing. DNA extraction, sequencing, and processing were carried out using the methods described in Bombin et al. (Bombin et al., 2020), with the following modifications. SILVA 132 Qiime release database was used as the reference database for the taxonomic assignment. Alignments were filtered by QIIME’s *filter_alignment*.*py* script. Reads cumulative sum scaling (CSS) normalization for alpha and beta diversities were performed through QIIME 1.91 with metagenomeSeq 1.26.1(Paulson et al., 2013). The phylogenetic trees of ZOTUs were assembled using the default options of QIIME 1.9.1 with the FastTree program (Price et al., 2009).

### Statistical Analysis

#### Bacterial Diversity

Alpha diversity indices (species richness, Shannon and Simpson diversity indices) were calculated using R package vegan v.2.5-6 (Oksanen et al., 2009). In order to evaluate the effect of diet and treatment on alpha diversity indices, as well as pairwise difference in alpha diversities between groups of samples, we applied a linear analysis of variance model using the **aov** function (base R) and Fisher’s LSD post-hoc test using DescTools v.0.99.36 (Signorell et al., 2020, Team and DC, 2019). Weighted Unifrac distances were calculated by beta_diversity_through_plots.py script with R 3.6.1 and Vegan v2.4-2 package (Oksanen et al., 2009). Bray-Curtis and Weighted Unifrac distances between groups were compared with permutational analysis of variance using the **anosim** function in vegan v.2.5-6 with 999 permutations (Oksanen et al., 2009).

#### Bacterial abundance

The difference in abundance of bacterial taxa was evaluated with the Wilcoxon test as described in Bombin et al. (Bombin et al., 2020). In order to evaluate if the diets could serve as categorical predictors for classification of the larvae bacterial samples as well as to determine which of the bacterial taxa drive the differentiation of the diets, linear discriminant analysis was performed with the **lda** function in MASS v.7.3-51.4 (Venables and Ripley, 2002) at the phylum, class, order, family, and genus taxonomic levels for the 10 most abundant representatives of the taxa at each taxonomic level. The first and second linear discriminants were visualized with a ggplot2 v.3.2.1 (Wickham, 2016).

The correlation between bacterial abundances on each diet was evaluated with the Spearman rank correlation test as described in Bombin et al. (Bombin et al., 2020). The only exception was the analysis of sterilized larvae raised on PHSA6 due to the limitation of available samples (4). The correlations in this group were evaluated with linear regression model using the **lm** function (base R) and **glance** function from broom v.0.7.0 (Robinson, 2014). At the ZOTU level, we used a linear regression model using JMP v.15.

To match the sequencing data with measured larval phenotypes, we randomly assigned each phenotypic measurement within each larvae group (based on diet, treatment, and genetic line) to one of the three groups and found an average per group. The correlation coefficients and p values were calculated with a Spearman rank correlation test as described in Bombin et al. (Bombin et al., 2020). Since we were only able to obtain four sequencing samples for S larvae raised on the PHSA6 diet (an insufficient sample size for Spearman’s rank correlation test), we evaluated the correlations with generalized linear quasi-Poisson models with the **glm** and **anova** functions (base R). The quasi-Poisson model was selected as it fits over dispersed data better than a linear regression model (Ver Hoef and Boveng, 2007, Bombin et al., 2016).

#### Larval phenotypes modeling

Normality tests, data transformations, and statistical models for larval phenotypes were done with JMP Pro 15.0. Phenotype measurements were tested for normality with the Shapiro-Wilk test and an outlier box plot. All phenotypic measurements data except survival were transformed for normality (S1). The influence of a diet on phenotype development was evaluated using a standard least squares model with post hoc t-test pairwise comparisons (Bombin et al., 2020) (S1).

## RESULTS

### Nutritional modifications influence larval bacterial community alpha and beta diversity

We were interested in whether addition of fat or sugar would lead to shifts in the bacterial community of the larvae. High dietary fat led to a significant increase in alpha diversity, assessed by Shannon’s index and species richness measurements, on both the regular and autoclaved-peach diets, but only in NS larvae (S2). Conversely, addition of sugar to the R diet caused a reduction in Shannon’s index (S2). Addition of fat or sugar was also associated with significant differences in beta diversity, as assessed by Bray-Curtis and weighted UniFrac distances, on both the regular and peach diets (Fig. 1, Sup. Fig1, S3). However, this was only significant in sterilized larvae.

**Figure 1:**
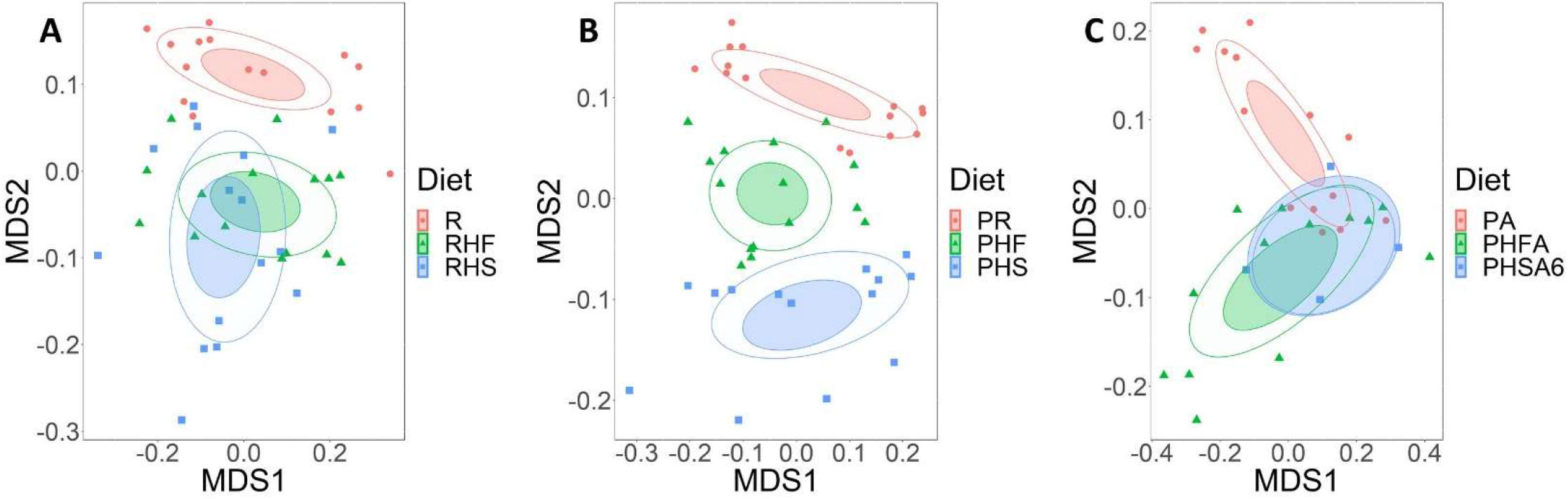
The response of bacterial community to dietary modifications varied with the origin of a diet. A) Sterilized larvae raised on a R diet formed a distinct bacterial community from larvae raised on RHF and RHS diets; B) Sterilized larvae raised on a PR diet formed a distinct bacterial community from larvae raised on PHF and PHS diets; C) Sterilized larvae raised on a PA diet formed a distinct bacterial community from larvae raised on PHFA but not PHSA diets. Bray-Curtis distances were calculated based on the abundance of all identified ZOTUs and visualized with a multidimensional scaling plot.

### Dominant bacteria taxa are associated with nutritional modifications

Using discriminant analysis, we evaluated which of the dominant bacteria taxa influenced the differentiation of the normal diet and its HF and HS modifications, at the level of family (Fig.2) and genus (Supp. Fig 2). NS larvae raised on the RHS diet were differentiated by *Enterobacteriaceae* and *Lactobacillaceae* at the family level (Fig. 2), and *Alsitipes* and *Blautia* at the genus level (Supp. Fig 2). The RHF raised larvae were discriminated by family *Bacteroidaceae* and *Corynebacteriaceae* (Fig. 2), and genus *Lachnospiraceae NK4A136* group (Supp. Fig 2). The R diet raised larvae communities were intermediate between the RHF and the RHS diets and primarily influenced by family *Acetobacteraceae* (Fig. 2) and genus *Pseudomonas* (Supp. Fig 2). The PHF and the PHS diets were the most separated from each other (Fig. 2). The PHS diet was differentiated by family *Enterobcteriaceae* and *Lachnospiraceae* (Fig. 2) and genus *Streptococus* (Supp. Fig 2). The PHF diet was separated by family *Erysipelotrichaceae* and *Corynebacteriaceae* (Fig. 2) and genus *Lachnospiraceae NK4A136* and *Alistipes* (Supp. Fig 2). The PR diet community was influenced by family *Ruminococcaceae* (Fig. 2) and genus *Pseudomonas* and *Gluconobacter* (Supp. Fig 2). The PHFA diet was differentiated by *Ruminococcaceae* and *Corynebacteriaceae* (Fig. 2), and genus *Gluconobacter* (Supp. Fig 2). The PA bacterial community was differentiated by *Lachnospiraceae* (Fig. 2) and genus *Pseudomonas* (Supp. Fig 2). The PHSA6 diet was not clearly separated from the PHSA11 and the PA diets’ communities at the family level, but was mostly influenced by *Lactobacillacelae*, and separated from PHSA11 by genus *Acetobacter* and *Lactobacillus*, while PHSA11 was discriminated by *Erysipelotrichaceae* (Fig. 2, Supp. Fig 2).

**Figure 2:**
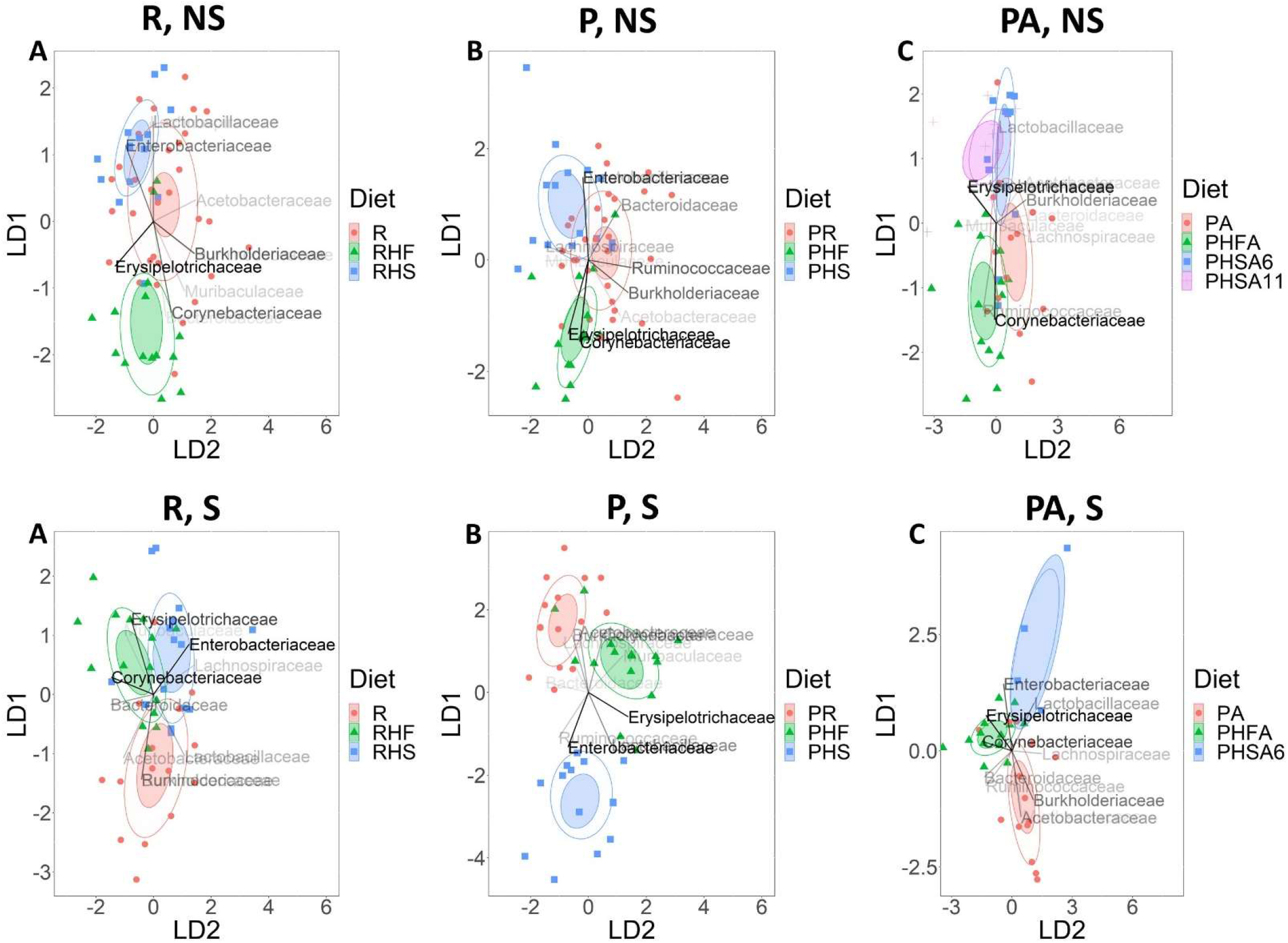
Nutritional modifications cause shifts in bacterial composition and are associated with changes in abundance of the dominant bacteria families. Linear discriminant analysis of 10 dominant bacterial taxa. A) Non-sterilized larvae raised on lab-based diets; B) Non-sterilized larvae raised on peach-based diets; C) Non-sterilized larvae raised on autoclaved peach-based diets; D) Sterilized larvae raised on lab-based diets; E) Sterilized larvae raised on peach-based diets; F) Sterilized larvae raised on autoclaved peach-based diets. The intensity of vector rays’ color corresponds to the strength of the impact that it produced on the samples to be separated in the vector direction, on a canonical plot. Confidence ellipses are filled based on the color of the diet. Normal data ellipses are unfilled and leveled to include 50% of the samples.

S larvae raised on the lab food-based diets were primarily defined by *Ruminococcaceae* and *Burkholderiaceae* (Fig. 2) and genus *Acetobacter* (Supp. Fig 2). The RHF diet was differentiated by *Erysipelotrichaceae* and *Muribaculaceae* (Fig. 2) and genera *Alistipes* and *Streptococcus* (Supp. Fig 2). The RHS was separated by *Enterobactericeae* and *Lachnospiraceae* (Fig. 2) and genus *Blautia* (Supp. Fig 2). The PR diet was differentiated by *Lachnospiraceae, Burkholderiaceae*, and *Acetobacteraceae* (Fig. 2), and genus *Acetobacter* (Supp. Fig 2). The PHF food raised larvae were separated by *Muribaculaceae* and *Corynebacteriaceae* (Fig. 2) and genus *Corynebacterium 1* (Supp. Fig 2). The PHS diet was defined by *Ruminococcaceae, Enterobacteriaceae*, and *Lactobacillaceae* (Fig. 2), and genera *Lactobacillus* and *Gluconobacter*. For the autoclaved diets, PA was defined by *Acetobacteraceae, Muribaculaceae*, and *Burkholderiaceae* (Fig. 2), and genera *Streptococcus* and *Acetobacter* (Supp. Fig 2). The PHFA diet was defined by *Corynebacteriacea* and *Erysipelotrichaceae* (Fig. 2), and genera *Corynebacterium 1* and *Lachnospiraceae NK4A136* (Supp. Fig 2). The PHSA6 diet was separated by *Enterobacteriaceae* and *Lactobacillaceae* (Fig. 2), and genera *Lactobacillus* and *Alistipes* (Supp. Fig 2). Furthermore, dominant bacteria taxa were influential in separating normal and modified diets at all taxonomic ranks (Supp. Fig 3-5).

**Figure 3:**
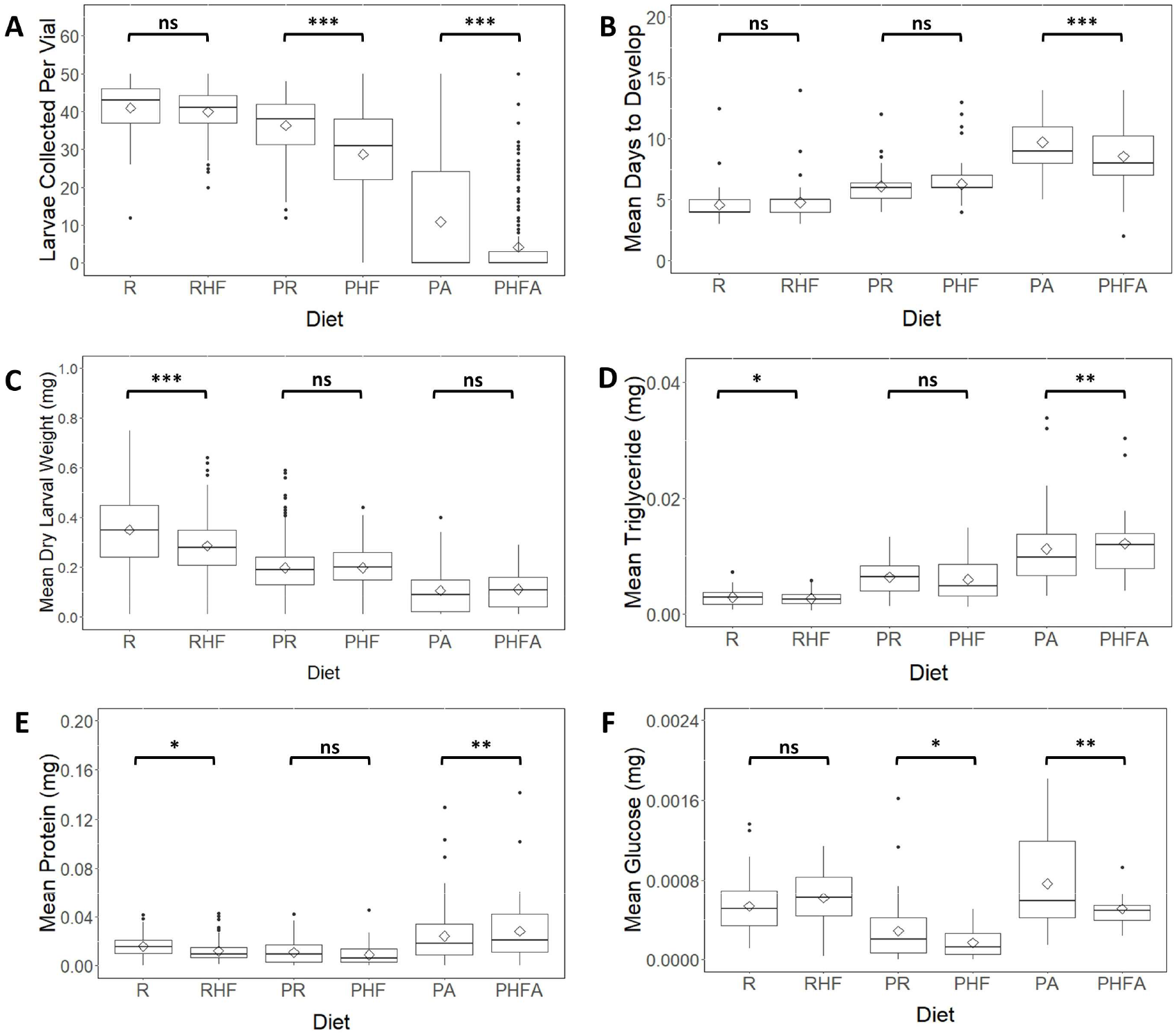
The response to high fat modification varies between sterilized larvae raised on peach-based and lab-based diets for most of the measured phenotypes. A) HF modification caused a decrease in mean survival until the 3^rd^ instar stage on peach-based diets; B) HF modification caused a decrease in larval development time only on a peach-based autoclaved diet; C) HF modification caused a decrease in larval weights on a lab-based diet only; D) HF modification caused an increase in larval triglyceride concentrations only on lab-based and autoclaved peach-based diets; E) HF modification caused a decrease in larval protein concentrations on a lab-based diet but an increase in protein levels on an autoclaved peach-based diet; F) HF modification caused a decrease in larval glucose concentrations only on the peach-based diets. Crossbars indicate the median, and rhombus indicate the mean. Asterisks indicate the significance of comparisons p < 0.001 ***, p < 0.01 **, p < 0.05 *, and ns p ≥ 0.05.

### Larval response to nutritional modification varies between lab-based and peach-based diets

On a lab diet, addition of extra fats increased larval triglyceride and glucose concentrations but decreased larval weight and protein concentration (Fig. 3, Fig. 4, S4). On a peach diet, the presence of additional lipids decreased larval survival, weight, and glucose concentrations (Fig. 3, Fig. 4, S4). The addition of extra fat to the PA diet decreased survival and development time (Fig. 3, Fig. 4, S4). In addition, sterilized larvae raised on the PHFA diet exhibited higher triglyceride and protein concentrations, but lower glucose concentration compared with the PA food raised larvae (Fig. 3, Fig. 4, S4).

**Figure 4:**
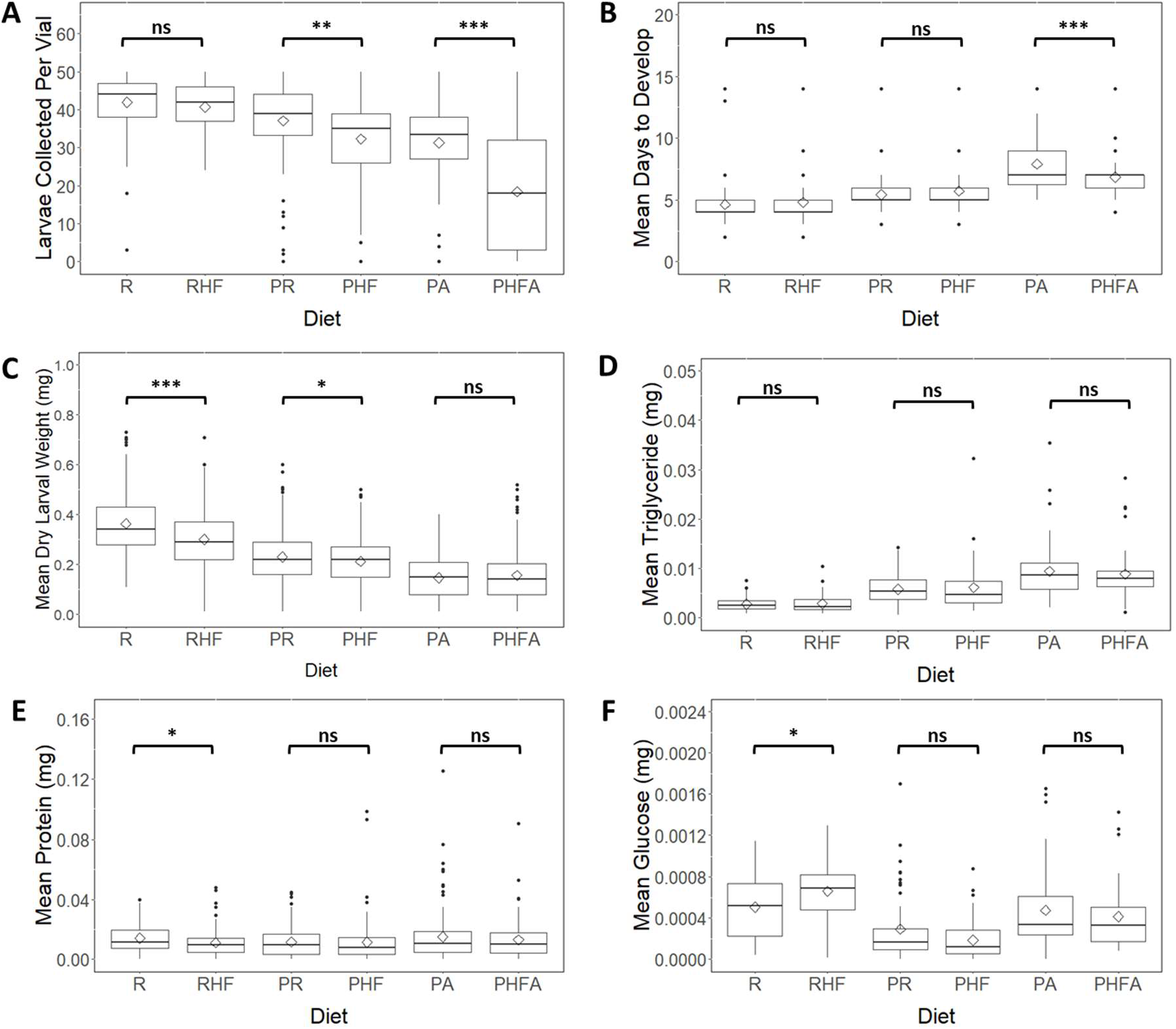
The response of non-sterilized larvae to high fat modification varies with a diet for most of the tested phenotypes. A) HF modification caused a decrease in mean survival until the 3rd instar stage on peach based diets; B) HF modification caused a decrease in larval development time only on a peach autoclaved diet; C) HF modification caused a decrease in larval weights on lab and regular peach diets; D) HF modification did not produce a significant effect on larval triglyceride levels; E) HF modification caused a decrease in larval protein concentrations only on a lab diet; F) HF modification caused an increase in larval glucose concentrations only on the lab diet. Asterisks indicate the significance of comparisons p < 0.001 ***, p < 0.01 **, p < 0.05 *, and ns p ≥ 0.05. Crossbars indicate the median, and rhombus indicate the mean.

Addition of extra sugar to the lab diet decreased larval weight (Fig. 5, Fig. 6, S4). In addition, it increased triglyceride concentrations for sterilized larvae and decreased protein levels for non-sterilized larvae (Fig. 5, Fig. 6, S4). HS modification of the peach diet resulted in a reduction of larval survival, longer development time, higher triglyceride and glucose concentrations (Fig. 5, Fig. 6). In addition, non-sterilized larvae raised on a PR diet were heavier than larvae raised on a PHS diet (Fig. 5, Fig. 6). Larvae raised on a PHSA6 diet developed faster, had higher weight, and glucose concentrations, but lower protein concentrations compared with PA raised larvae (Fig. 5, Fig. 6, S4). Interestingly, more sterilized larvae survived on the PA diet compared with PHSA6 diet, while opposite was true for non-sterilized larvae (Fig. 5, Fig. 6, S4). In addition, sterilized larvae raised on PHSA11 diet exhibited lower survival, development time, and protein concentrations, but higher weight (Fig. 5, Fig. 6, S4). Non-sterilized larvae raised on PHSA11 diet had longer development time, higher glucose concentration, and lower protein concentration, when compared with PA raised larvae (Fig. 5, Fig. 6, S4).

**Figure 5:**
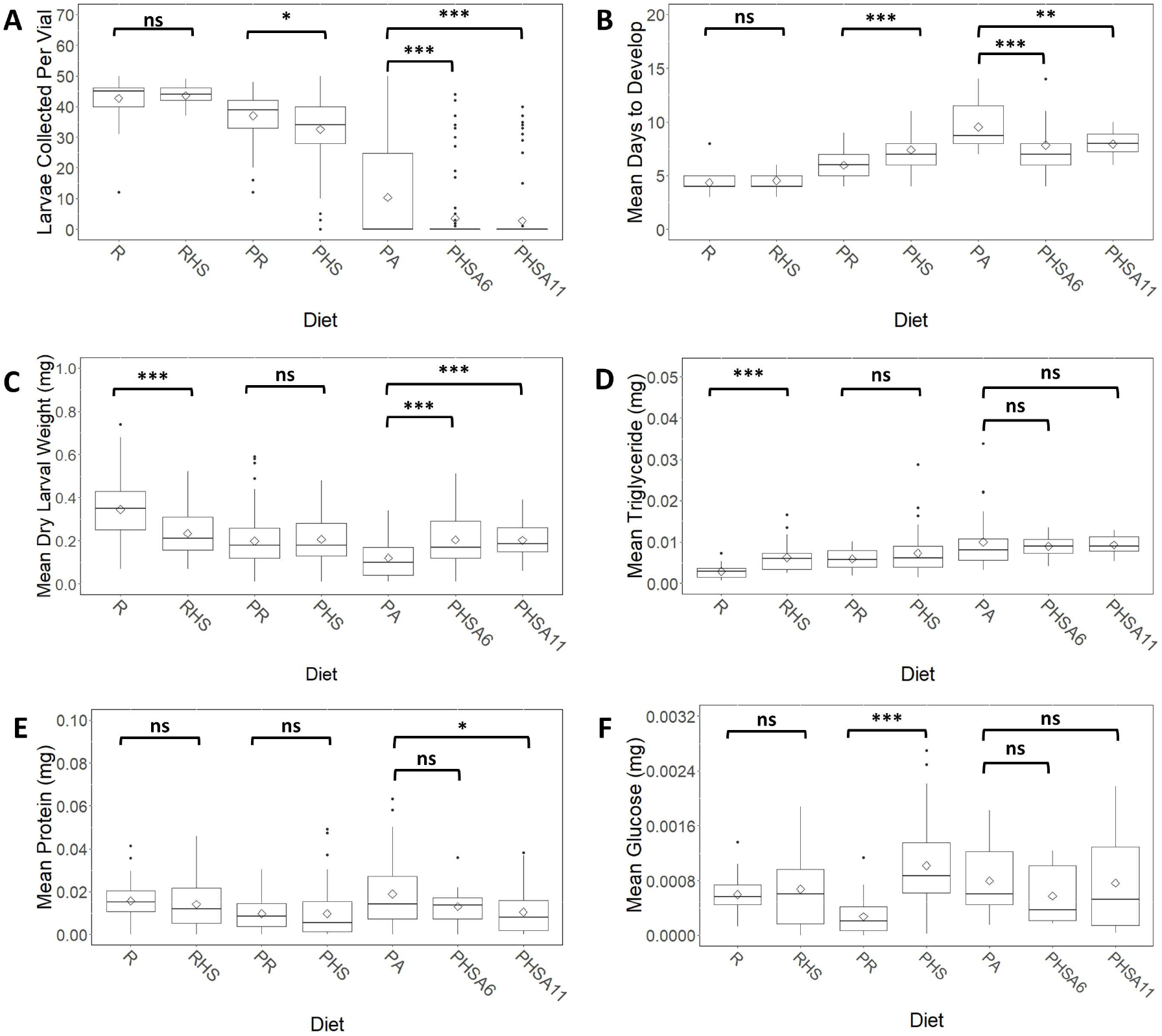
Sterilized larvae raised on lab-based and peach-based diet exhibit a different response to high sugar nutritional modification, for most of the measured phenotypes. A) HS modification caused a decrease in mean survival until the 3^rd^ instar stage on peach-based diets; B) HS modification caused an increase in development time only on a peach-based diet; C) HS modification caused a decrease in larval weights only on a lab-based diet; D) HS modification caused an increase in triglyceride concentrations only on a lab-based diet; E) HS modification caused a decrease in larval protein concentrations only on an autoclaved peach-based diet; F) HS modification caused an increase in larval glucose concentrations only on a peach-based diet. Crossbars indicate the median, and rhombus indicate the mean. Asterisks indicate the significance of comparisons p < 0.001 ***, p < 0.01 **, p < 0.05 *, and ns p ≥ 0.05.

**Figure 6:**
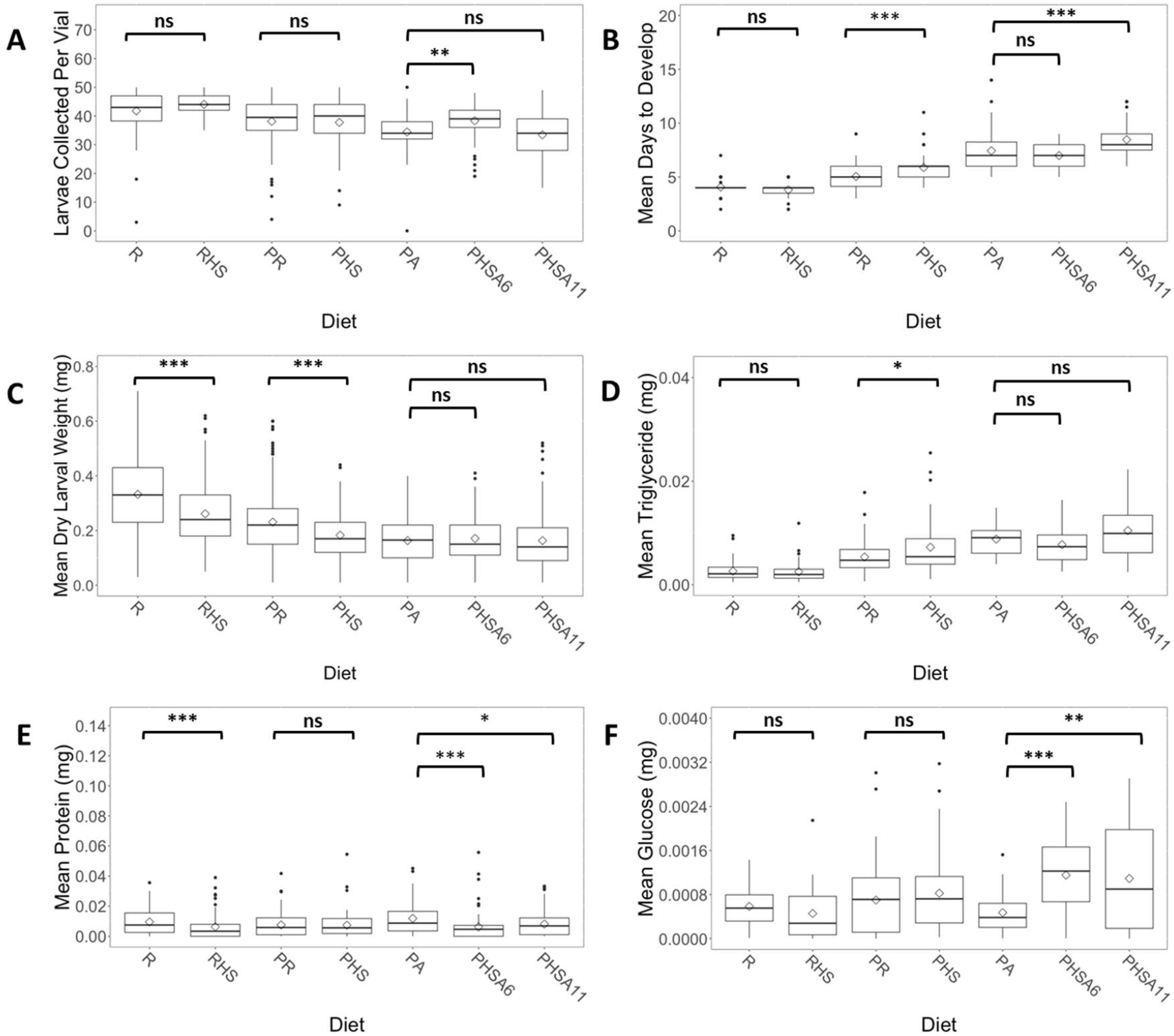
The response of non-sterilized larvae to high sugar modification varies with a diet. A) HS modification increased larval survival only on a PHSA6 diet; B) HS modification increased larval development time on most of the peach-based diets. C) HS modification decreased larval weight on a lab and peach diets; D) HS modification increased larval triglyceride levels only on a peach diet; E) Protein concentrations of only larvae that were raised on a peach diet did not change significantly due to HS modification; F) Glucose concentrations of the larvae raised on PA diet significantly increased due to HS modification of the diet. Crossbars indicate the median, and rhombus indicate the mean. Asterisks indicate the significance of comparisons p < 0.001 ***, p < 0.01 **, p < 0.05 *, and ns p ≥ 0.05.

## DICSCUSSION

### Bacterial community composition varies with larval diets

A very limited number of studies that compared *Drosophila* gut bacterial communities in response to HF or HS modifications of the diet are currently available. Von Frieling et. al (2020) suggested that HF bacterial community is distinct from the normal diet. Jehrke et. al (Jehrke et al., 2018) did not observe substantial differences in the structure of the whole bacterial communities of normal and HS diets. In our work, largely, standard and modified diets exhibited a significant difference in the community structure and phylogenetic diveristy as was indicated by Bray-Curtis and Unifrac distances, especially if larvae were sterilized. Comparing normal and modified diets, we observed that HF diets were more similar in their taxonomic diversity to normal diets than were HS diets, particularly for sterilized larvae. Considering vertebrate models, previous research indicated that mice and rats fed on HF and HS diets formed bacterial communities that were distinct from ones feeding on a normal diet (von Frieling et al., 2020, Do et al., 2018). As described above, we observed similar patterns in our data, especially for sterilized larvae, suggesting that macronutrient content is an important determinant of bacterial community structure.

### Abundances of the dominant bacteria taxa are associated with diet types

Among the dominant bacteria groups, we also noticed several taxa that were consistently associated with certain diet modifications. HF diets were associated with Actinobacteria and Bacteroidetes at the phylum level, Corynebacteriales and Erysipelotrichales at the order level, *Corynebacteriaceae* and *Erysipelotrichaceae* at the family level, and *Corynebacterium 1* and *Lachnospiraceae NK4A136* group at the genera level. Several representatives of *Corynebacterium* are known to be members of symbiotic microbiota of vertebrates, exhibit lipophilic qualities, and produce lipolytic enzymes (Hahne et al., 2018). In our work, *Corynebacteriaceae* was negatively correlated with the glucose concentrations on RHF and PHF diets. *Corynebacterium 1* was negatively correlated with glucose concentrations, on the same diets. *Corynebacterium* (a different taxon from *Corinabacterium 1*) was positively correlated with glucose on PHFA and negatively correlated with glucose on PHF diet. Previous research indicated an increase in the abundance of *Erysipelotrichaceae* in mice fed a high fat diet (Kaakoush, 2015). *Erysipelotrichaceae* was also shown to be associated with host lipid metabolism and dyslipidemic phenotype (Martínez et al., 2013). In our work, we observed positive correlations between *Erysipelotrichaceae* and triglyceride concentrations and the development rate on PHF diet and glucose concentrations on PHFA diet. *Lachnospiraceae* was also shown to be correlated with high-fat diets, altered lipid metabolism and obesity in humans and animal models (Vacca et al., 2020). In our work, *Lachnospiraceae* was positively correlated with glucose concentrations on PHFA diet.

HS diets were associated with Tenericutes and Fermicutes at the phylum level, Lactobacillales and Enterobacteriales at the order level, *Enterobacteriaceae* and *Lactobacillaceae* at the family level, and *Lactobacillus* at the genus. *Lactobacillus* is known to produce glycoside hydrolases and polysaccharide lyases which can degrade carbohydrates (Wang et al., 2020). Previous research showed that *Lactobacillaceae* abundances were increased in honeybees that were fed a sucrose solution; the same carbohydrate that we used in order to make HS diets (Wang et al., 2020). *Lactobacillus* strains are known to be a part of human microbiota and were correlated with a decrease in fasting glucose sugar and insulin resistance (Azad et al., 2018, Khalili et al., 2019). Mice that were fed by high sucrose diet were shown to have higher abundances of *Lactobacillus* than mice fed on a normal diet (Magnusson et al., 2015). The survival of ingested *Lactobacillus plantarum* was shown to be improved upon feeding on high fat high sugar diets (Yin et al., 2017). In our work, on HS diets, we observed that *Lactobacillaceae* was positively correlated with total number of larvae and development rate on PHS diet and negatively correlated with protein and triglyceride concentrations on RHS diet. *Lactobacillus* was positively correlated with the total number of larvae and development rate on PHS diet. In addition, a negative correlation was observed with weight on a PHS diet and protein and triglyceride concentrations on a RHS diet. *Enterobacteriaceae* and other members of Protobacteria are known for their ability to utilize simple carbohydrates rapidly and for being associated with a high sugar diet (Satokari, 2020, Volynets et al., 2017, Park et al., 2013, Magnusson et al., 2015). In addition, *Enterobacteriaceae* were shown to be correlated with the obese phenotype (Xiao et al., 2014). In our work, *Enterobacteriaceae* was negatively correlated with triglyceride concentrations on PHS diet, positively correlated with development rate on PHS and larvae survival on PHS and PHSA6 diets. These findings suggest that several symbiotic taxa have similar associations with multiple host organisms. Therefore, these relations might be evolutionarily conserved.

The regular lab diet was also consistently associated with certain bacterial taxa, such as Virrumicrobia and Epsilonobacteria at the phylum level, Pseudomanales and Acetobacterales at the order level, *Acetobacteraceae* and *Burkholderiaceae* at the family level, and *Pseudomonas* and *Acetobacter* (mostly for sterilized larvae). Members of *Acetobacteraceae* family are known to be a part of normal *Drosophila* microbiota in the lab and wild flies (Vandehoef et al., 2020, Han et al., 2017, Bost et al., 2018, Winans et al., 2017). In our previous work, we observed that on unmodified diets, *Acetobacteraceae* was negatively correlated with triglyceride concentrations on the PA and the PR diets, positively correlated with weight on the PA and on the PR diets and positively correlated with development time on the PR diet (Bombin et al., 2020). Moreover, *Acetobacter* was positive correlated with weight on the PA and the PR diets and development time on the PR diet(Bombin et al., 2020). Several members of *Burkholderriaceae* and *Pseudomonas* are known to be pathogens in *Drosophila* (Vodovar et al., 2005, Pilátová and Dionne, 2012, D’Argenio et al., 2001). Our previous work found a negative correlation between *Burkolderriaceae* and weight on the PR diet and larvae development time on the R diet (Bombin et al., 2020). *Pseudomonas* was negatively correlated with the total number of larvae on R diet as well as development time and weight on a PR diet (Bombin et al., 2020). Triglyceride concentrations on PR diet and development time on PA diet were positively correlated with the abundance of *Pseudomonas* (Bombin et al., 2020).

### Symbiotic microbiota may alleviate some of the negative effects produced by harmful dietary modifications

On the modified peach diets, presence of symbiotic microbiota reduced the mortality and development time of the larvae. This is consistent with previous research showing that the parental microbiota helps larvae to adapt to nutritionally unfavorable conditions (Bing et al., 2018, Berger et al., 2005, Henry et al., 2020b). Similarly, high pre-adult mortality of flies and increased development time was shown on a high sugar low-yeast diet, especially when symbiotic microbiota was removed (Wong et al., 2014, Henry et al., 2020b).

Consistent with our previous work, diets that supported higher development and survival rates also generally supported higher weights and lower triglyceride concentrations of the larvae (Bombin et al., 2020). It was shown that an increase in sugar and fat concentrations may cause a decrease in larvae weights and increase in triglyceride concentrations (Reed et al., 2010, Musselman et al., 2011, Jehrke et al., 2018, Galenza et al., 2016, Wong et al., 2015). In our work, presence of environmental and parental microbiota often reduced the negative effect of diet modification on larval weight and triglyceride concentrations. However, sterilized larvae raised on autoclaved peach diets, especially with harmful modifications exhibited extremely low survival rate and mostly were sampled from two genetic lines. That complicated the evaluation of a general pattern for metabolic phenotypes and suggested a presence of genetic adaptations in those *Drosophila* lines.

Consistent with our previous findings, the glucose and protein concentrations in larvae did not directly correlate with life history traits (Bombin et al., 2020). Previous research indicated that a high sugar diet may cause increased glucose concentrations in *Drosophila* and vertebrate models (Galenza et al., 2016, Do et al., 2018, Jehrke et al., 2018, Wong et al., 2014). We found similar results for sterilized larvae raised on peach-based diet and for non-sterilized larvae raised on autoclaved peach-based diets. Although Galenza et. al (2016) and Jehrke et. al (2018) reported no significant correlations between increased sugars consumption and protein concentrations; on the natural diet, we observed that presence of symbiotic microbiota can alleviate a reduction in larval protein concentration caused by the high sugar diet. In addition, our results indicated that presence of symbiotic microbiota reduces the difference in glucose and protein concentrations caused by addiction of extra lipids to a peach diet.

### Conclusions

Consumption of high fat and high sugar diets is often associated with a development of unhealthy metabolic phenotype and obesity (Seganfredo et al., 2017, Patterson et al., 2016, Al-Goblan et al., 2014). We observed that nutritional modifications of the diets are associated with shifts in gut bacterial community composition. Interestingly, the dominant bacteria taxa associated with particular nutritional modification (high fat or high sugar), in our *Drosophila* model are similar with bacterial community members associated with westernized diet, in vertebrate models, and human populations. Microbiota acquired from the environment or inherited maternally are capable of reducing negative metabolic effects, caused by nutritional modification of a diet, especially on peach-based diets.

