## Supplement figures for "Symbiotic bacterial community of *Drosophila melanogaster* changes with nutritional modifications of the diet but can alleviate negative effects on larval phenotypes"

### Slide 1
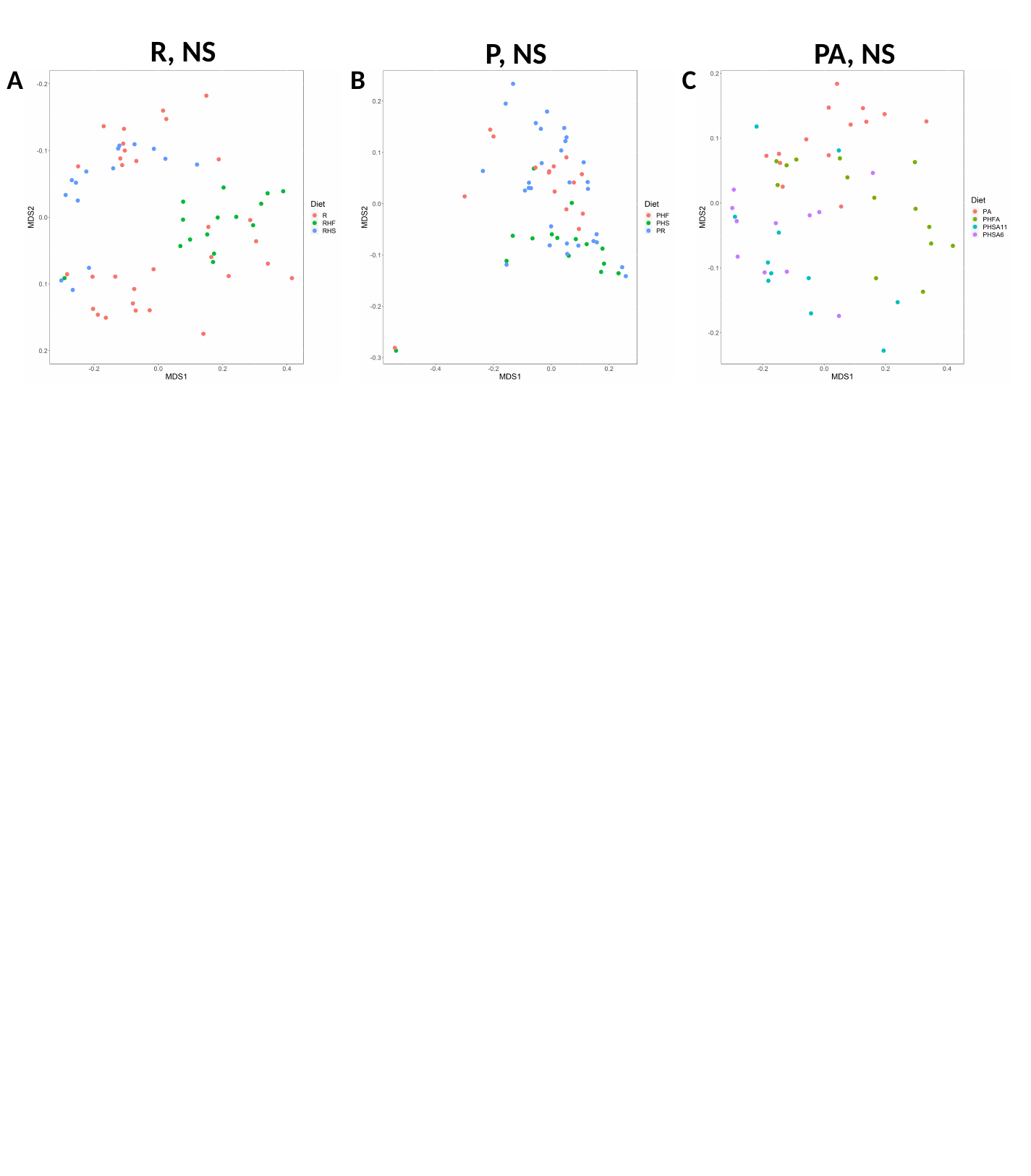

R, NS
P, NS
PA, NS
A
B
C

### Slide 2
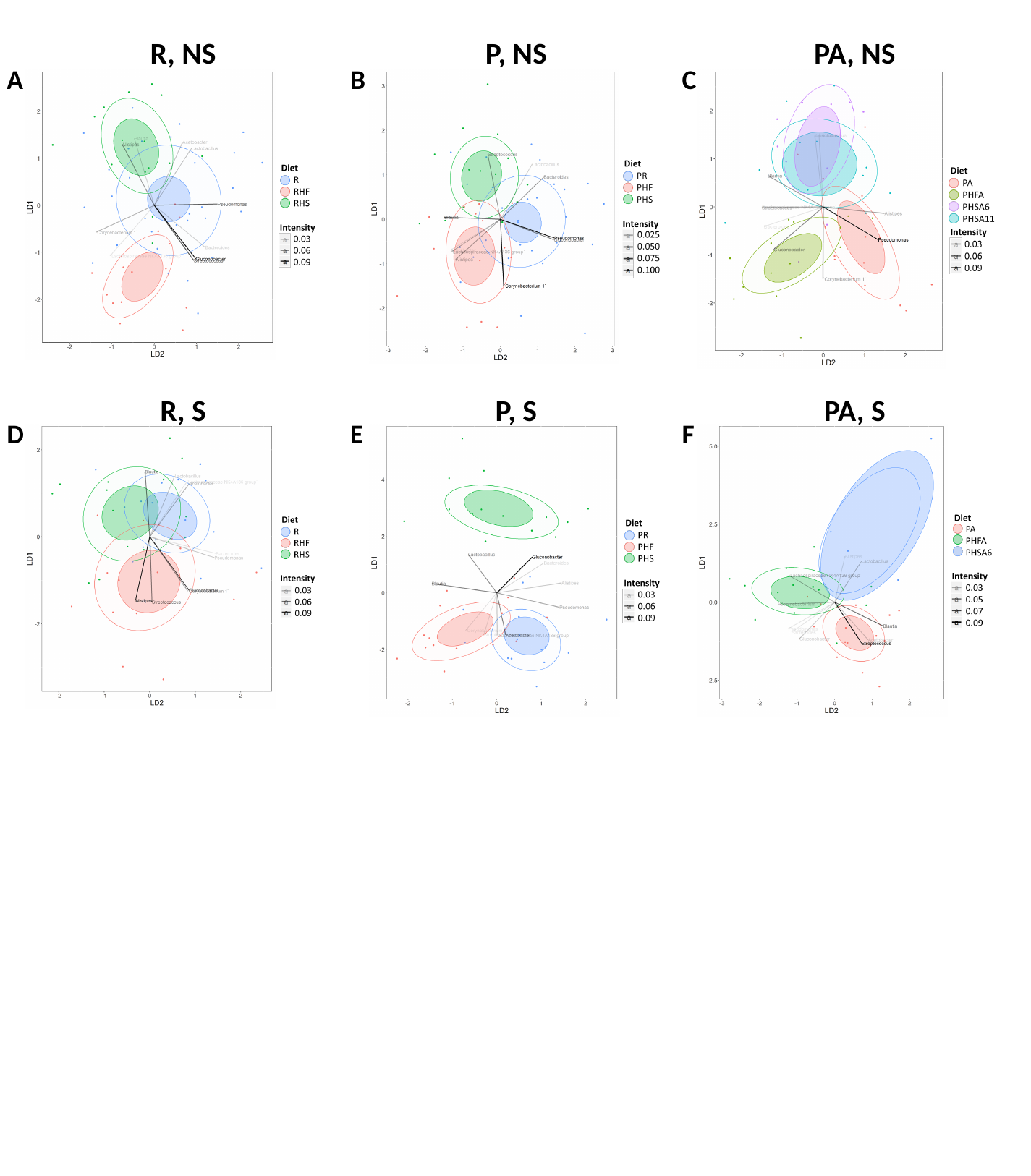

R, NS
P, NS
PA, NS
A
B
C
R, S
P, S
PA, S
D
E
F

### Slide 3
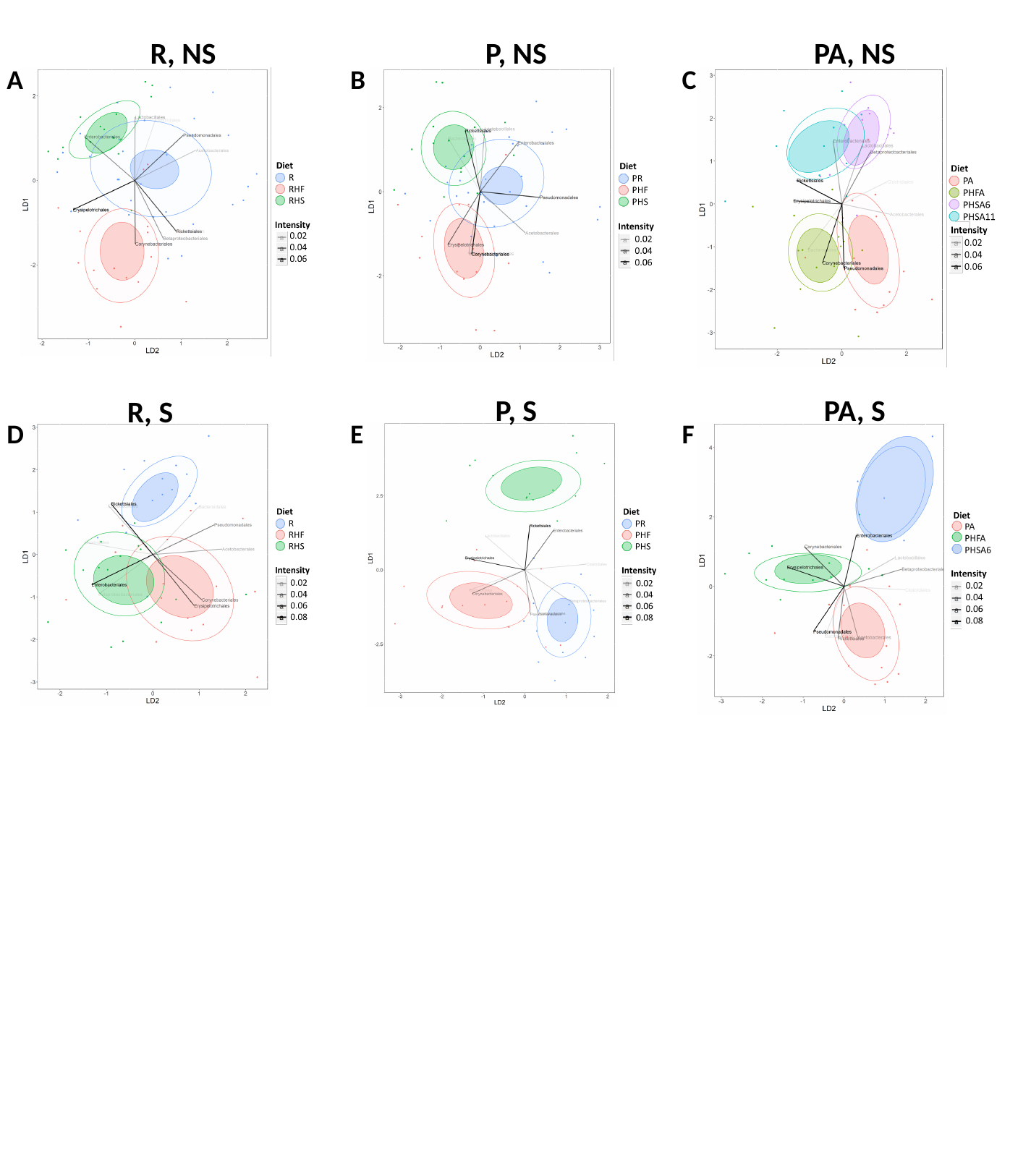

R, NS
P, NS
PA, NS
A
B
C
P, S
PA, S
R, S
D
E
F

### Slide 4
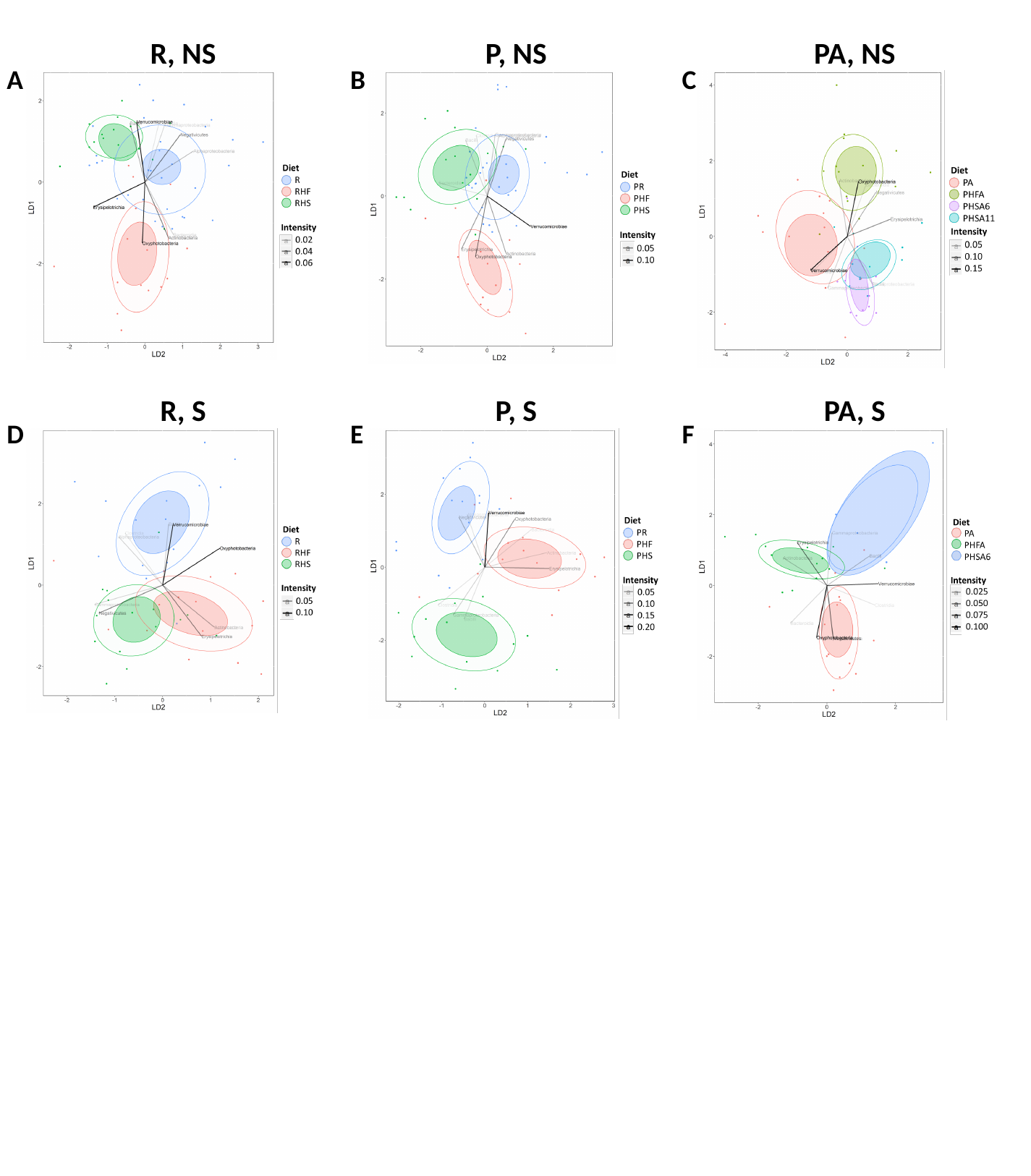

R, NS
P, NS
PA, NS
A
B
C
R, S
P, S
PA, S
D
E
F

### Slide 5
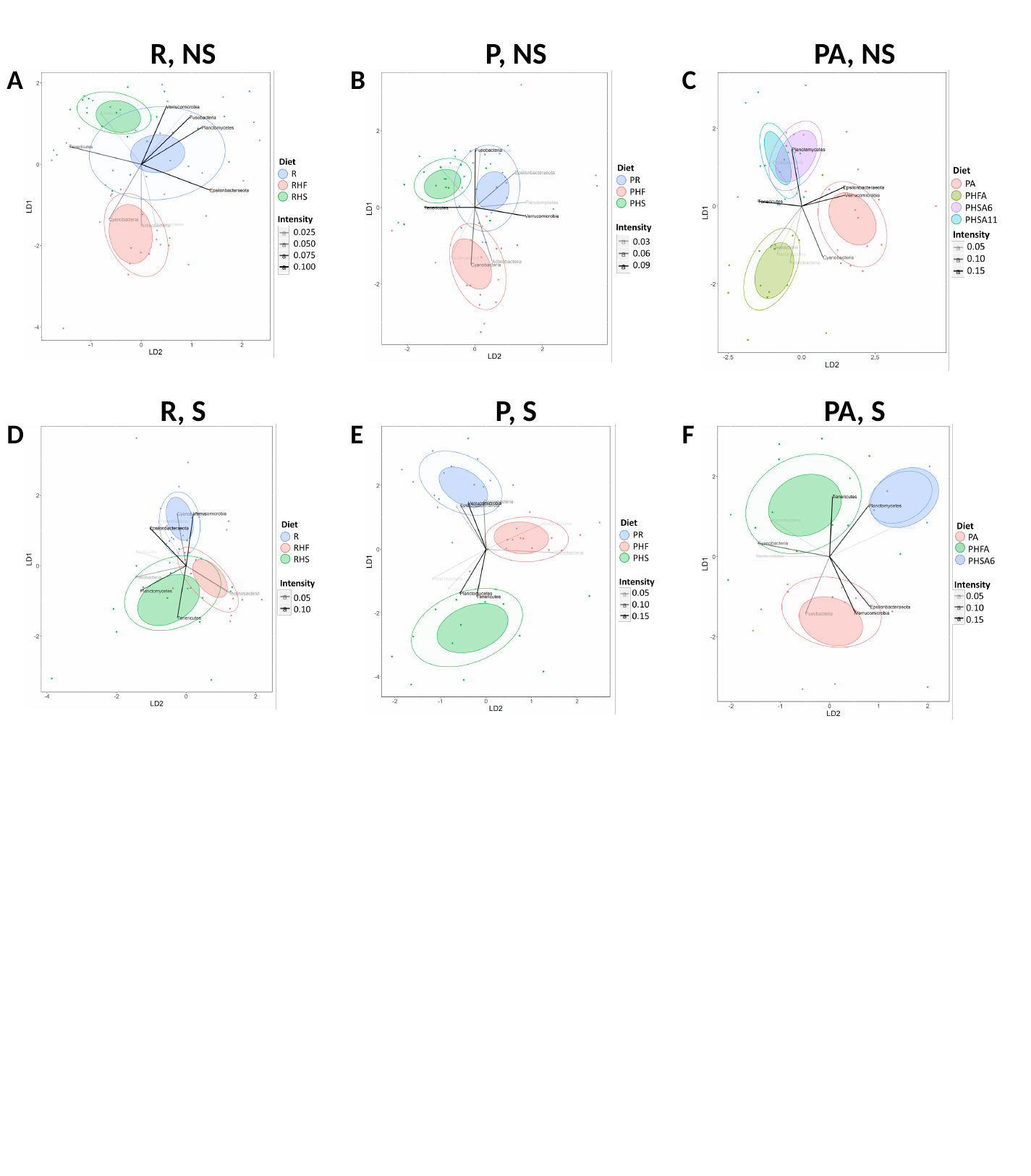

R, NS
P, NS
PA, NS
A
B
C
R, S
P, S
PA, S
D
E
F
